# Similarities Between Somatosensory Cortical Responses Induced via Natural Touch and Microstimulation in the Ventral Posterior Lateral Thalamus in Macaques

**DOI:** 10.1101/2021.11.10.468076

**Authors:** Joseph Thachil Francis, Anna Rozenboym, Lee von Kraus, Shaohua Xu, Pratik Chhatbar, Mulugeta Semework, Emerson Hawley, John Chapin

## Abstract

Lost sensations, such as touch, could be restored by microstimulation (MiSt) along the sensory neural substrate. Such neuroprosthetic sensory information can be used as feedback from an invasive brain-machine interface (BMI) to control a robotic arm/hand, such that tactile and proprioceptive feedback from the sensorized robotic arm/hand is directly given to the BMI user. Microstimulation in the human somatosensory thalamus (Vc) has been shown to produce somatosensory perceptions. However, until recently, systematic methods for using thalamic stimulation to evoke naturalistic touch perceptions were lacking. We have recently presented rigorous methods for determining a mapping between ventral posterior lateral thalamus (VPL) MiSt, and neural responses in the somatosensory cortex (S1), in a rodent model (Choi et al., 2016; Choi and Francis, 2018). Our technique minimizes the difference between S1 neural responses induced by natural sensory stimuli and those generated via VPL MiSt. Our goal is to develop systems that know what MiSt will produce a given neural response and possibly a more natural “sensation.” To date, our optimization has been conducted in the rodent model and simulations. Here we present data from simple non-optimized thalamic MiSt during peri-operative experiments, where we MiSt in the VPL of macaques with a somatosensory system more like humans. We implanted arrays of microelectrodes across the hand area of the macaque S1 cortex as well as in the VPL thalamus. Multi and single-unit recordings were used to compare cortical responses to natural touch and thalamic MiSt in the anesthetized state. Post stimulus time histograms were highly correlated between the VPL MiSt and natural touch modalities, adding support to the use of VPL MiSt towards producing a somatosensory neuroprosthesis in humans.

## Introduction

Our overall aim in this line of work is to find a method that would allow us to use MiSt or other neural stimulation modalities, and emulate natural neural responses in the somatosensory cortices (S1) and other somatosensory regions as determined necessary for the perception of touch. We hypothesize that similarity of neural responses following MiSt and tactile stimulation will translate into similarity of perceptions. On the other hand, MiSt that produces “unnatural” neural patterns will not result in “natural” touch perception. Therefore, if we can determine the best locations and patterns to produce such naturalistic neural responses, we should create more natural sensations. We may need to consider the neural response in a more extensive set of structures to fine-tune this approach to achieve this goal. Here, we present our findings in a non-human primate (NHP) model. We started with responses in S1 to natural touch as our template in which to optimize our VPL MiSt-induced responses as shown in our previous rat and simulation-based studies (Brockmeier et al., 2011; Choi et al., 2012, 2016; Li et al., 2013b, 2013a, 2015; Choi and Francis, 2018). We note that the results presented in this paper were recorded circa 2008 and the above optimization methods had not been developed or implemented. However, we feel these results from the NHPs should be shared as we move towards human implementation of these systems, where we can directly interrogate our underlying hypothesis that more naturalistic S1 responses lead to more naturalistic sensations. The rodent somatosensory system is significantly different from humans and NHPs we utilized (Francis et al., 2008). Here we used simple non-optimized MiSt in the acute NHP preparation and show that the somatotopy is generally well-maintained with VPL MiSt, comparable to natural touch in the NHP, as in the rodent (see discussion).

It has been demonstrated that neuronal activity in the motor cortex can be used to directly control computer cursors and robotic systems via a BMI (Chapin et al., 1999; Serruya et al., 2002; Taylor et al., 2002; Hochberg et al., 2006; Ganguly and Carmena, 2009; Ajiboye et al., 2017; Degenhart et al., 2020). Recently, interest in BMIs has exploded as it has become clear that such systems are likely to restore motor function lost due to spinal cord injury (SCI), neurological disease, or amputation. Such BMIs require a closed-loop configuration that uses not only real-time neural data to move an actuator, such as a robotic arm, but also delivers sensory feedback to the user (Flesher et al., 2021). To date, this feedback has come mainly through the intact visual system of the user who is viewing their performance with the BMI. However, it is known that natural reaching and dexterous tasks require somatosensory feedback for high levels of performance and control. Therefore, somatosensory feedback from sensors on a neurally controlled prosthetic arm/hand presented directly to the user via MiSt of the neural substrate should lead to a better controlled prosthetic. This somatosensory feedback, along with visual feedback, helps control such devices and allows them to become one with the user.

The use of cortical MiSt to directly introduce information into the brain has been demonstrated with some success (Talwar et al., 2002; Fitzsimmons et al., 2007; London et al., 2008; O’Doherty et al., 2011; Flesher et al., 2016, 2021). Several investigators have shown that macro- and microelectrode stimulation in the human somatosensory thalamus (ventral caudal nucleus Vc, referred to as the Ventral Posterior Lateral in other mammals (VPL), and nearby thalamus) can produce a variety of somatosensations, including both natural and artificial, ranging from perceptions of touch or movement to sensations of hot or cold, tingling, or the sense of pressure (Lenz et al., 1995; Davis et al., 1998, 1999; Kiss et al., 2003b; Ohara et al., 2004; Chien et al., 2017). In many cases, the elicited sensation depended on the stimulus frequency and its amplitude (Patel et al., 2006). Proprioceptive and cutaneous sensory modalities were found to segregate between different regions of the thalamus as described in the literature (Sacco et al., 1987; Kaas, 2007; Francis et al., 2008). This separation should help produce separate touch and proprioceptive channels for sensory input via MiSt or other stimulation modalities.

Although human thalamic studies have been beneficial in demonstrating conscious perceptions induced by (Vc/VPL) electrical stimulation, we still lack a method for producing reliably “normal” sensations. When conducting intraoperative experiments on humans, there are several constraints, such as the amount of time one must rigorously explore the stimulation state space and ethical concerns that limit the areas from which one can sample. Much work on humans has utilized large (>1 mm) macroelectrodes, which are often intended for deep brain stimulation (DBS) to alleviate movement disorders such as tremor or to alleviate chronic pain. This stimulation is generally at high frequencies (100-300 Hz). Kiss et al. (Kiss et al., 2003a) have suggested that such macrostimulation may activate a neural area 4000-fold greater than MiSt. This more extensive activation may cause tingling or other paresthetic sensations through the widespread recruitment of axons and neurons.

With these limitations in mind, as an initial step toward developing an optimized somatosensory neuroprosthesis in humans, we have utilized multielectrode neurophysiological techniques in encephalated NHPs (Macaca radiata) to determine how thalamic MiSt might be used to evoke neural responses in the somatosensory cortex (S1). As the macaque has a similar somatosensory stream to humans, it is a more suitable animal model for such work as compared to the rodent model (Kaas, 2007; Francis et al., 2008). Our protocol involved implantation of multi-electrode arrays in the hand representations of the VPL (VPL; 4 electrodes) and primary somatosensory (S1) cortex (32 electrodes). These simultaneous recordings allowed us to record the response patterns of hundreds of single and multi-units in the S1 cortex during computer-controlled natural touch stimulation and MiSt in somatotopographically equivalent areas of the VPL. These acute experiments allowed us to directly and quantitatively compare the post-stimulus responses evoked by natural touch and VPL MiSt in S1 in anesthetized subjects intraoperatively. We used MiSt in the low-frequency range (≤ 5 Hz), with the work presented here held to just one biphasic MiSt pulse in the VPL. Here we show that simple MiSt in the VPL elicits S1 cortical responses with similar properties, such as amplitude and duration, to those induced via natural touch, at least in the anesthetized state. Thus, we add evidence that utilizing such VPL-MiSt may be suitable for a somatosensory neuroprosthesis.

## METHODS

All work adhered to NIH Guide for the Care and Use of Laboratory Animals and was approved by SUNY Downstate’s IACUC and followed the recommendations of the Weatherall report, “The use of non-human primates in research”. All efforts were made to minimize animal suffering, including anesthetics for all surgical procedures (see below). For this work, animals were given ad lib food and water.

### Surgical preparation and recordings

All experiments were conducted in the acute anesthetized preparation. Monkeys (Macaca radiata) were initially anesthetized with Ketamine (15mg/kg) and intubated to allow controlled ventilation and administration of Isofluorane at 0.5-3% in (95% O_2_). Fentanyl (I.V.) 2-5 mcg/kg/hr was used throughout the surgery to further ensure no pain would be felt. After anesthesia had been established, the animal was placed in a stereotactic frame. A midline incision was made, and the skin retracted above the central sulcus. Craniotomies were performed directly over the arm regions in S1 and above the VPL thalamic nucleus (see atlas (Paxinos et al., 2000)). We first implanted the S1 cortical electrode arrays in the granular layers and then performed a series of natural touch stimulation experiments to define the precise skin-to-cortex representation in that animal. The implanted cortical electrodes were left in place for the remainder of that experiment.

Next, we slowly drove the thalamic electrode array into the VPL thalamus guided by the macaque atlas (Paxinos et al., 2000). We stopped at 100 μm intervals to record neuronal RFs and then MiSt at different currents while simultaneously recording multi and single-unit activity from the S1 electrode array. We obtained large data sets that provided a comprehensive record of the somatotopographic relationships between the skin, the VPL, and the S1 cortex. Subsequently, we searched a subset of the VPL stimulus parameter space. To maintain consistency of the subjects for a given experiment, the NHPs hand was held in place with a ring stand and flexible cord. At the same time, the tactor was attached to a second ring stand and positioned to touch the desired region of skin.

We will refer to the brain region stimulated by the touch of a single point on the periphery as a **Stimulus Field**(**SF**) compared to a **Receptive Field**(**RF**), which is the peripheral domain that can elicit a response from a single neuron. Specifically, the region of S1 that is stimulated by either touch of a point on the periphery, an S1-SF, or to VPL-MiSt at a point in the VPL that is a VPL-MiSt induced S1-SF.

### Electrodes

Implanted electrodes consisted of an array of 32 sharp (35 μm diameter) Tungsten electrodes for acute NHP experiments. Each array was arranged in 2 parallel rows, each with 16 electrodes spaced ~300 μm apart. After craniotomy and removal of the dura to expose the hand area in the vicinity of the central sulcus, the electrode array was positioned on the lip of the post-central gyrus such that the anterior row of electrodes was placed about 1 mm caudal to the central sulcus and its parallel row was placed 1 mm caudal to that. This puts the electrode array in area 1 on the cortical surface but into area 3b as one drives the array deeper (Paxinos et al., 2000). We have pooled our data in much of the analysis here and thus do not make claims to be recording from area 1 or 3b specifically.

All the electrode tips were placed flush on the cortex and then slowly driven down until layers III-IV were reached. These electrodes then remained in place, allowing the same single and multiunit clusters to be recorded simultaneously during up to 165 stimulation experiments. Our VPL array consisted of 4 sharp stainless-steel electrodes in a square array with 1.0 mm separation. This array was used for both recording and stimulation. The VPL array was progressively driven down through the VPL’s somatotopic representation of the cutaneous periphery, allowing us to record and stimulate for a series of experiments.

### Neural recording and analysis

The Plexon Inc. multichannel acquisition processor (MAP) system was used for online data acquisition and spike discrimination. An offline sorter was used for post-hoc re-discrimination. The Plexon Offline Sorter provided a variety of methods for post-hoc single unit discrimination. Conventional approaches were used for general spike separation and removal of stimulus artifacts. Data analysis utilized the NEX neurophysiology analysis system and its Matlab and Excel extensions. Statistica was used for statistical analysis and plotting.

We simultaneously recorded multiple neuronal waveforms from each of the electrodes and then performed offline discrimination. First, we lumped together the neural recordings from each electrode, allowing accurate estimation of the neural population responses from each cortical location. We then used a peak detection algorithm to find the maximum response from all electrodes. The simplest and most reliable method was to record the multi-unit activity from each electrode, use computer algorithms to measure the maximal response peaks and background activity in post-stimulus histograms, and then convert the response amplitudes into Z-scores, which could be normalized across the entire electrode array.

To minimize contamination from the VPL MiSt artifact, we blanked out the first 2 ms following stimulation, which should be under the amount of time it takes for conduction of an action potential from the VPL to the S1 cortex as well as subsequent action potential generation in the S1 cortical recipient neurons. In addition, we sorted the stimulus artifact as a unit in our template sorting method stated above and did not include these “units” representing the stimulus artifacts, which are very stereotypical and easily clustered, in our analysis.

### Tactile stimuli

Mechanical touch stimuli were applied to different regions of the hand and forearm using a computer-driven vibromechanical actuator to deliver mechanical pulses to the skin. Our standard stimulus was a single pulse producing <1 mm skin displacement for ~1ms, delivered at rates of 5 Hz. The touch experiments involved serially tapping up to 12 locations on the hand. These results were then compared with electrical MiSt in the VPL. A single experiment lasted for 90 seconds and included tapping at only one position on the hand. Likewise, all microstimulation experiments lasted for 90 seconds and included stimulating in one electrode configuration with a given stimulus waveform.

### Multichannel microstimulator

We developed a modular 16-channel bipolar constant-current MiSt system capable of producing any arbitrary pattern of brain stimuli through multi-electrodes. Single and/or 2-electrodes were employed to produce MiSt. All VPL stimuli were made using bipolar stimuli to minimize the stimulus artifact through closely spaced pairs of electrodes. All stimuli were biphasic, normally with the anode first. Cathode-first trials were also investigated but did not produce obvious differences. MiSt pulse widths ranged from 100-500 μs. Stimulus currents ranged from 25-100 μA. Stimuli consisted of a single biphasic pulse. For all the data presented in this paper, the MiSt was biphasic and bipolar utilizing 200 μsec duration phases. We did not specifically search the MiSt state space for exact thresholds; instead, we used 25, 50, 75, and 100 μA as our test amplitudes. These amplitudes were chosen after brief preliminary work that spanned responses from “weak” to “strong” and enveloped the natural touch responses, as can be seen in the figures.

## RESULTS

A total of 357 recording experiments were conducted on 3 NHPs. All utilized simultaneous recordings from the S1 cortical hand area using spaced electrode arrays consisting of 2 rows of electrodes (2×16 electrodes). These experiments yielded neural activity from stimulus fields driven by a natural touch of the hand, or MiSt in the VPL at varying stimulus intensities, where a stimulus field is defined as the brain region responsive to touch at a single point on the periphery, or MiSt at a single point in the VPL. Each experiment typically involved approximately 450 stimulus presentations (at 5 Hz) of touch or VPL MiSt. All NHPs also received a 2 × 2 electrode array implanted in the VPL thalamus.

### Qualitative results

In Fig.1, we have plotted the raw post-event-rasters and their associated post-event-time-histograms below them induced via a natural touch of the fingers. For plots with numbers less than 8 (e.g., sig001i), these are multiunit activity recorded from the VPL, and channels above 8 (e.g., sig021i) are from the S1 cortex. Notice the variety of responses, some with early phasic response, others with a later phasic response, and some with both an early and a late response such as sig001i. In Fig.2, we have plotted the same type of activity as in Fig.1, but for the MiSt-induced responses in the cortex. In Fig.2, all the raster histogram pairs are in response to 25μA biphasic stimulation. In addition, for two multi-units, we have plotted the responses to several amplitudes of MiSt as denoted in the key. Note the different scales on the y-axis. Most of these units that had a significant response also had a simple phasic response, as seen in Fig.1.

**Fig 1.**
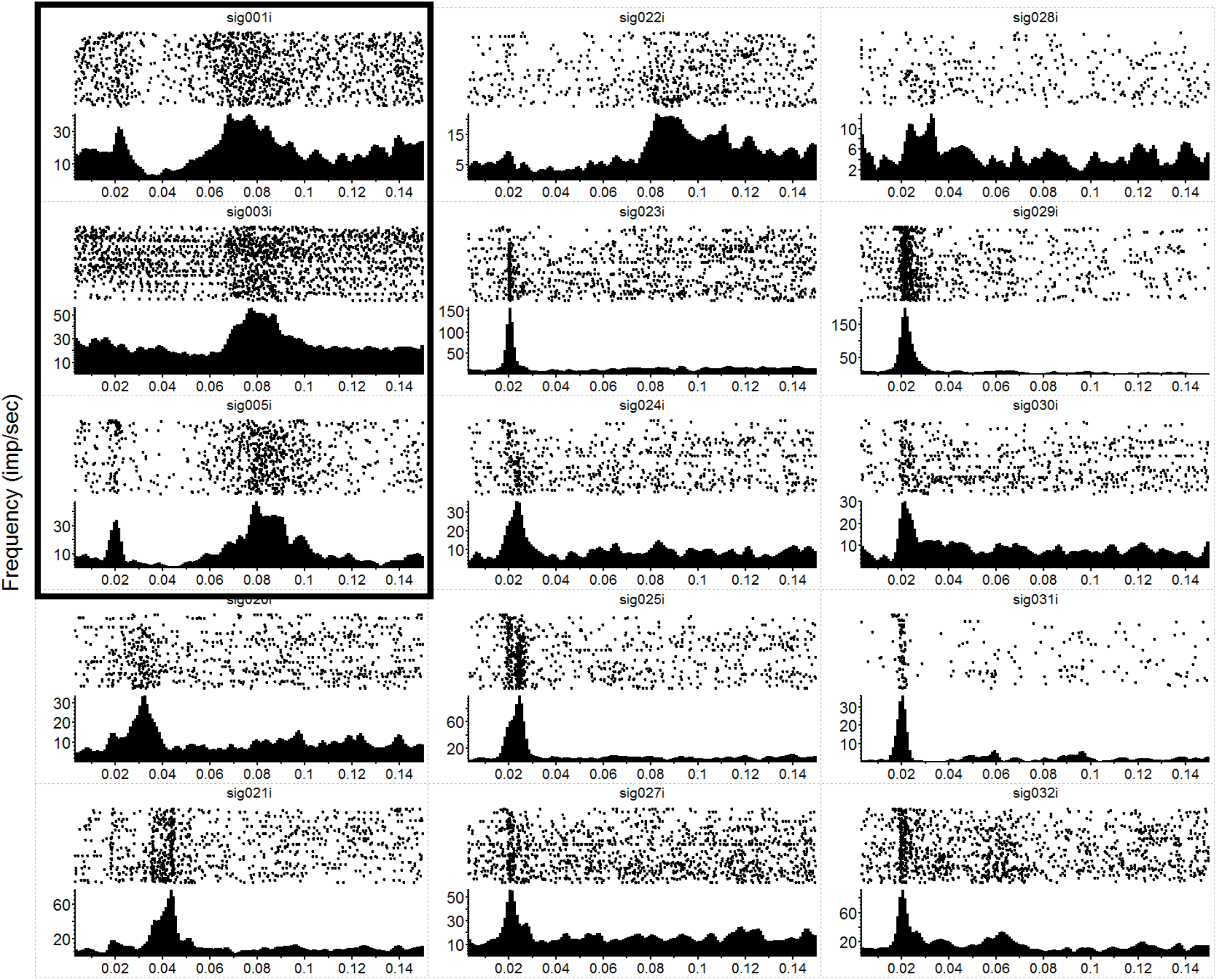
Neural Responses to Mechanical Stimulation. Raw data showing Post-Stimulus-Time-Rasters with their corresponding Post-Stimulus-Time-Histograms below for a subset of the 36 recording electrodes during a single mechanical touch experiment. In this experiment, we touched the anesthetized NHP’s hand at 1 position at a frequency of 5 Hz. Each raster-histogram pair is labeled with the electrode channel number, where sig < 8 are VPL thalamic channels (first 3 panels on the left column, marked by the surrounding box) and sig > 8 are S1 cortical channels. The i indicates that these are unsorted units; thus, we are showing the multiunit activity recorded on each channel. There was a 4-electrode array in the thalamus (2 × 2, with 1mm spacing) and a 32-channel array in the cortex (2 × 16, with an intra-row spacing of 300 μm and inter-row distance of 1 mm). Note the diversity of responses and the differences in the y-axis, which is the unsorted units firing rate in Hz, while the x-axis is time in seconds. PSTH bins were 1 ms and smoothed with a 3 ms Gaussian moving window. The time axis starts at 3 ms to match the x-axis with Fig. 2 (where MiSt stimulus artifact required blanking of the first couple of ms).

**Fig. 2.**
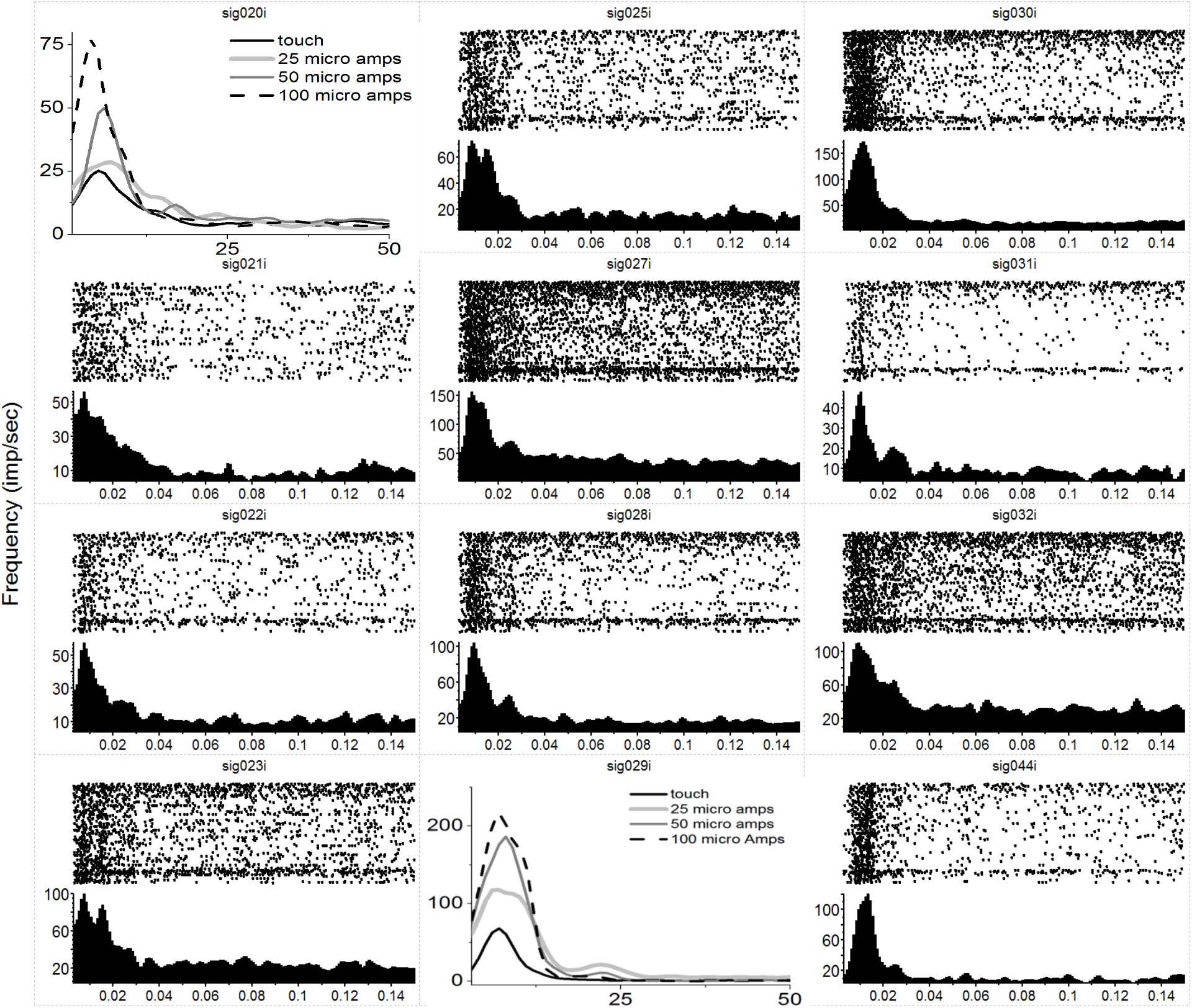
Neural Responses in S1 to MiSt in the VPL thalamus. Each peri-event raster and histogram pair are in response to a 25μA single biphasic bipolar pulse stimuli in the VPL. For panels showing multiple histograms, the current used is labeled in the key, and responses from the mechanical touch were shifted in time to align with the MiSt-induce responses; the x-axis for these plots is bin number at 3ms bins. Processing was as in Fig. 1. Note that most responses are on the order of 15 – 20 ms.

**Fig. 3.**
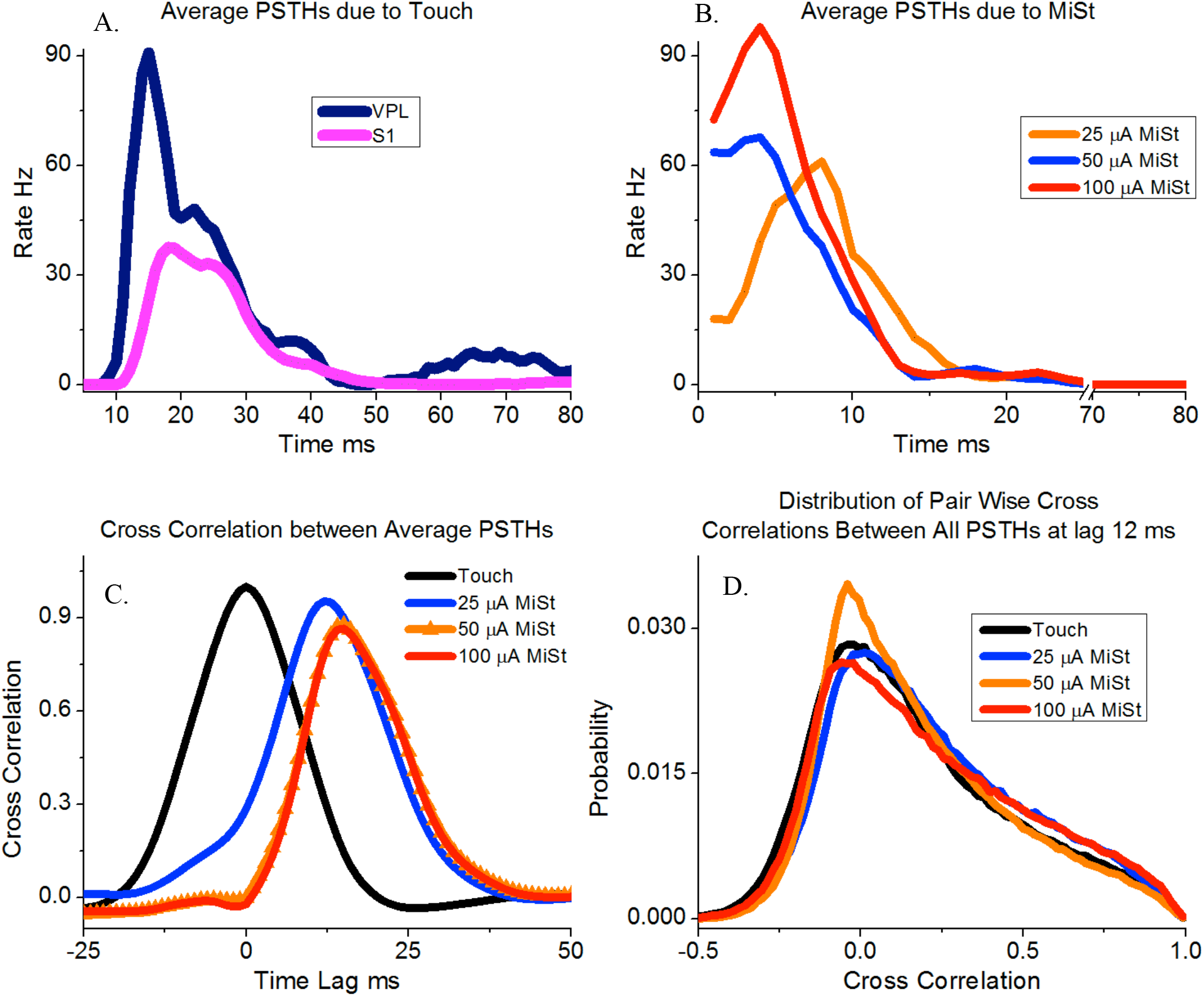
**A**, Average thalamic and S1 neural responses induced by the touch of the periphery using only channels that had a significant response for this average (N = 1239). **B**, Responses for three amplitudes of microstimulation, again only using the channels with significant responses defined as peeks of 3 STDs or more (N = 438, 631 and 420 for 25, 50, and 100 μA respectively). **C**, Crosscorrelation between the average PSTHs for touch vs. the three MiSt amplitudes used most during our study. **D**, Pair-wise crosscorrelation distribution between all pairs of PSTHs between touch and MiSt amplitudes. The touch-induced PSTHs were time-shifted by 12 ms as this led to the smallest sum of squared differences between the three MiSt distributions and the touch-induced distribution.

As this was in response to MiSt, the responses are all shifted to the left; they occur sooner than would be the case if due to touch on the periphery as there is no peripheral transmission delay.

### Somatotopy

We found an obvious relationship (somatotopy) from the peripheral touch to the VPL as expected from the literature on these and other mammals (Krubitzer and Kaas, 1992; Kaas, 2007; Francis et al., 2008). In addition, we witnessed the expected somatotopy between the VPL and primary somatosensory cortex (Kaas, 2007) and found the VPL MiSt maintained these relations. Thus, VPL MiSt on an electrode responding to digit one touch would produce comparable S1 responses in the same cortical area activated by a natural touch of digit one.

The results from a set of typical experiments on an anesthetized NHP are shown in Fig. 4, where we have drawn a cartoon of the NHP hand color-coded according to induced neural responses found in either the VPL (Fig.4.C) or S1 (Fig.4.B). Panel B depicts the electrode array in S1 color-coded based on points of the hand and their associated S1-SF, with VPL-SF shown in panel C. In panel D, we show the bipolar pairs of electrodes that were used for VPL-MiSt. Note that VPL electrodes 1 and 2 had strong responses to touch (SF) while electrodes 3 and 4 had weak responses, implying that 3 and 4 were just outside the core VPL. In support of this is the fact that all pairs lead to just one S1-SF except for the strong response pair of VPL electrodes (1 and 2), which leads to a response of both S1-SFs that are concordant with those SFs seen in the thalamus. Thus, these results imply that if we have thalamic receptive fields for each of the digits as well as those tessellating the palm, we should be able to generate S1 cortical responses representing any portion of the hand.

**Fig. 4:**
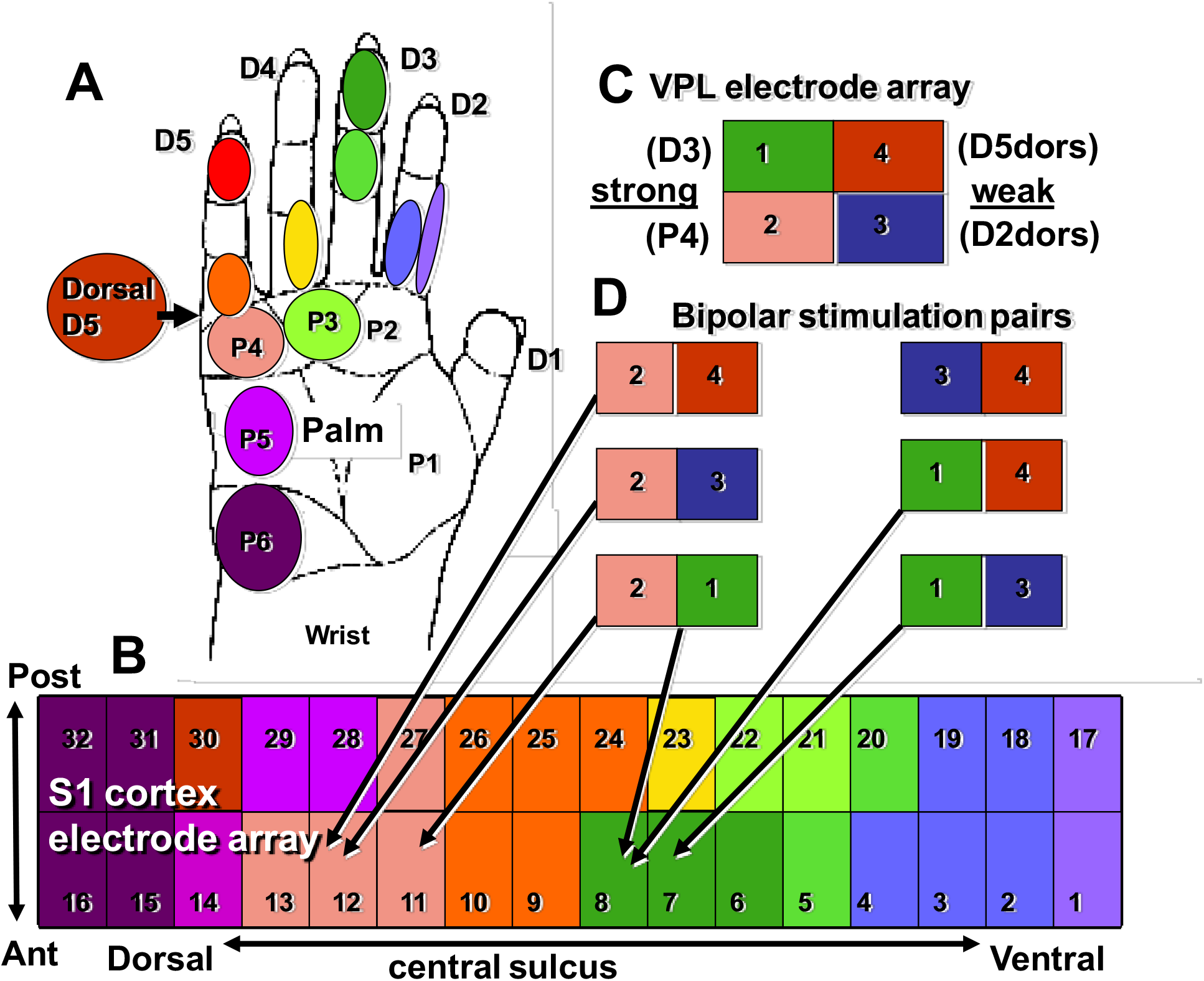
This cartoon Test results for concordance between responses in the somatosensory cortex to natural touch and VPL-MiSt through electrodes with similar stimulus fields to those seen on the S1 electrodes. **A**. Diagram of the NHP’s hand labeled and color-coded with positions that were stimulated via our tactor. These same colors are used to describe the stimulus fields on the S1 electrodes (**B**) and VPL electrodes (**C**). **D**. Are the VPL electrode pairs used for our bipolar microstimulation. Note that two of the four VPL electrodes recorded strong SF to touch (1, 2) while the other two were weak (3, 4) and possibly on the boundary of the VPL. The terms D2dors and D5dors are the digit number on the dorsal surface. Due to this arrangement, the cortical stimulus fields to VPL stimulation are governed by the strong VPL channels. See Choi et al. 2016 for a similar relationship in the rat.

In Fig. 5, we describe the topographic associations between the periphery, the VPL, and the S1 cortex. Our electrode array was situated caudal to and parallel with the central sulcus. Electrodes 1 and 17 were most lateral and electrodes 16 and 32 most medial, the peak induced neural activity forms a diagonal band as seen in Fig.5. This banding simply reaffirms the known somatotopy. For instance, it is known that digit one is represented lateral to digit 4, which can be seen in this figure as channel 1 has a peak in activity in the lateral electrodes numbered one and 17. In contrast, the peak activity for digit four is seen more medially on electrodes 3, 4, 5, 18, 19, and 21. As stated the color code at the right depicts the z-score for each point, e.g. 2 = p < .05 and 3 = p < .003. One can see in Fig.5 that some of the tactile stimulation-induced responses are focal such as for D1, whereas others are more diffuse such as for P4. As all the MiSt in these experiments were bipolar, with electrodes separated by 1 mm, the neural responses could have two peaks, such as for the <u>D4-P1,2</u> stimulation, where underlining indicates this was a VPL-MiSt induced response.

**Fig. 5:**
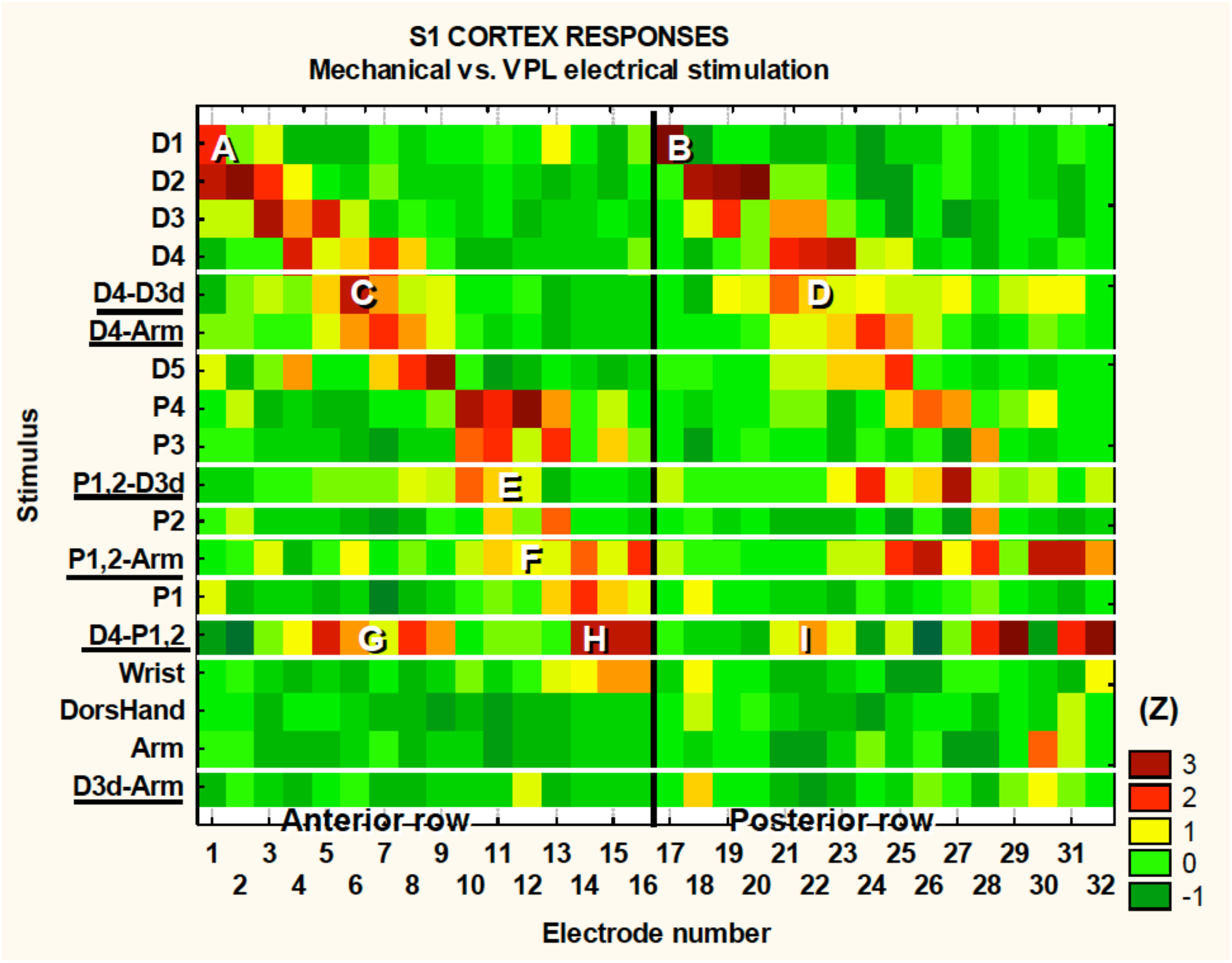
S1 somatotopy in response to touch and VPL-MiSt. Shown are the results from 18 SFs recorded in S1 during 12 natural touch and 8 VPL-MiSt experiments (underlined, e.g., <u>D4-Arm</u>). VPL-MiSt was between two electrodes, where one electrode could have one SF, such as D4, and the other could be an Arm SF. Thus, the VPL-MiSt is labeled by both (D4-Arm). We have ordered the data in the expected somatotopic progression starting with D1. The color code at right depicts the z-scored significance for each point, e.g., 2 = p < .05 and 3 = p < .003 (see methods).

### MiSt Responses

In general, MiSt at currents up to 100 μA produced localized responses within S1. Fig.6.A shows examples of 4 averaged cortical responses sequentially recorded during natural touch of Digit3 and MiSt in the VPL - Digit3 representation at currents of 25, 50, and 100 μA. These results were typical in that both VPL MiSt and natural touch stimuli produced peaked cortical responses with a prominent center flanked by decaying surrounds. This basic pattern was consistent across our sample. Fig.6.B shows the average of 15 cortical responses recorded during natural touch experiments and 7 from 75 μA VPL MiSt. These averaged responses depict the statistical means and standard errors for each electrode in an S1 cortical array. We have aligned all the data such that the peaks correspond. These results demonstrate that: 1) VPL MiSt produces distinctly peaked cortical responses in S1. 2) The cortical response amplitudes and widths correlate with stimulus current (Fig.6.A). 3) The cortical responses induced via VPL MiSt are comparable to those induced via natural touch (Fig.6.B).

**Fig. 6:**
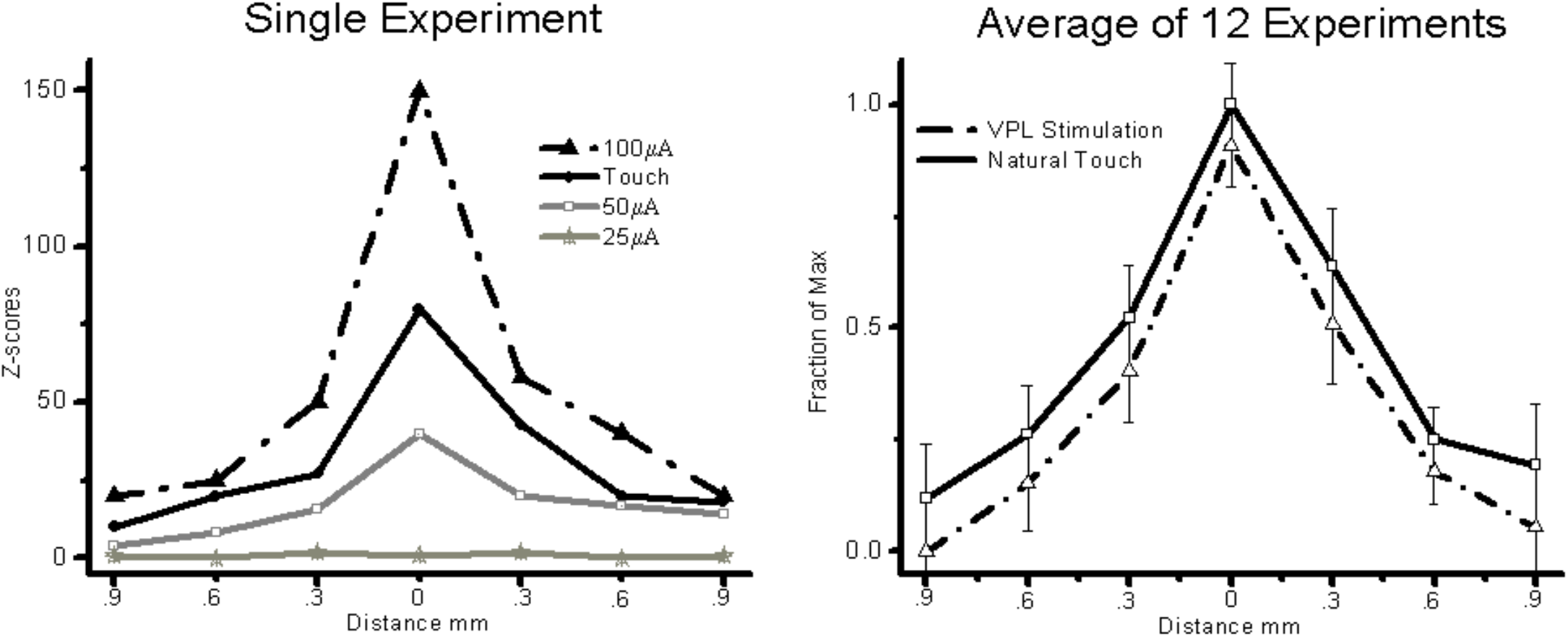
VPL MiSt produces SFs comparable to natural touch in S1. **A**, SFs produced via VPL-MiSt or touch from a typical experiment. **B**, Averaged stimulus fields from 15 natural touch and 7 bipolar VPL stimulation experiments. VPL-MiSt was at 75 μA. Each group depicts a pyramidal stimulus field ±standard errors (error bars). Note the similarity between the two curves. All MiSt were biphasic bipolar pulses with each phase 200 μsec long. The X-axis is the distance in mm between cortical electrodes.

## Discussion

In this paper, we have presented results that indicate MiSt in the VPL thalamus of NHPs can induce neural responses in S1 similar to those produced via natural touching of the hand, as we have previously demonstrated in the rat model (Choi et al., 2016; Choi and Francis, 2018). Despite the worry that VPL stimulation might provoke widespread nonspecific neural responses, we measured precise matches between the stimulus fields in the VPL stimulus sites and the stimulus fields in the activated regions of the S1 cortex. Small areas of single digits were readily discerned. This may be because the VPL thalamocortical fibers rapidly conduct somatotopically congruent axon bundles to circumscribed koniocortical target zones in S1. In contrast, antidromically activated corticothalamic fibers are relatively slow and dispersed. We observed that the sizes of the VPL-MiSt SFs in S1 cortex were tightly correlated with stimulus current, suggesting that the current spread approximately spherically in the thalamus before transmitting directly to the 2D surface of S1. We have conducted modeling of this spread in the rat (Choi et al., 2012), which improves our ability to model VPL MiSt and S1 activation patterns. Bipolar stimulation between two VPL electrodes spaced more than 1 mm apart produced two separate response areas in the S1 cortex. This suggests that the stimulation mainly occurred in the high-current density regions around the electrodes, which has been shown using optical techniques in other sensory areas (Histed et al., 2009).

The ability of the brief VPL MiSt to emulate the spatiotemporal characteristics of S1 cortical responses to simple natural touch implicates the thalamocortical path as the major determinant in at least initiating these responses, as would be expected. On the other hand, direct cortical MiSt leads to widespread inhibition that typically lasts for 100 ms in the rat with only a very brief initial excitatory response lasting about 2 ms (Butovas and Schwarz, 2003). We found that strong VPL stimulation in NHPs produced 5 or more oscillatory responses in the S1 cortex, while natural stimulation generally only produced 2-3 oscillatory responses. These oscillations occur at approximately 600 Hz and have been discussed previously (Baker et al., 2003). These results suggest that a possible mechanism for paresthesias is the highly synchronous nature of the VPL stimulation. Our conjecture, therefore, is that the ideal VPL-MiSt is one that closely approximates the response patterns produced by natural stimuli not only in the spatial extent, which has been the focus of this report but also concerning the fine temporal structure of the responses, as accomplished in our rodent work (Choi et al., 2016). However, we ultimately need to move such work into humans to address questions on the qualia of the evoked sensations.

The work we have presented was conducted with an anesthetized NHP preparation. However, we have obtained similar results in the awake restful state in rats indicating these ideas will at least hold in that neural state (Brockmeier et al., 2011). It seems prudent to replicate this in the awake NHP before moving to humans. Indeed, this is just the beginning of this work, as we expect that the awake, actively sensing state would be more complicated. We may need to consider information about other brain regions that feed into S1, such as the motor cortices, in addition to the current state of the S1 cortex and VPL while utilizing VPL-MiSt. Recently it has been shown that S1 cortical activity is modulated by reward expectation (Pantoja et al., 2007; Ramakrishnan et al., 2017; Atique and Francis, 2021), punishment expectation (Yao et al., 2021) and there delivery. Indicating such affective modulation should also be tested with somatosensory neuroprosthesis.

Recently much work has been conducted using MiSt of S1 cortex directly to evoke percepts (Talwar et al., 2002; London et al., 2008; O’Doherty et al., 2011; Flesher et al., 2021). Work such as that conducted by Romo et al. has shown that S1 MiSt can be used by nonhuman primates as somatosensory information in the flutter vibration domain (Romo et al., 1998, 2000), while others have shown artificial proprioception via learning (Dadarlat et al., 2015). In addition, our previous work utilizing the Roborat rat paradigm has allowed us to demonstrate somatosensory neural prosthetic capabilities in a rat model. We used MiSt of the primary somatosensory cortex in this previous work with clear results (Talwar et al., 2002). Our preliminary results using this same paradigm with VPL-MiSt have been successful, indicating that at least in the rat, such VPL-MiSt with single biphasic stimuli, like those presented here, are perceivable by the animal (data not shown).

Our neuroprosthetic techniques utilizes the production of a neural template generated via the natural peripheral sensory organ, such as the skin for touch in our case, and working toward minimizing the difference between that template and cortical responses induced via MiSt, such as VPL-MiSt in the present case (Brockmeier et al., 2011; Li et al., 2013b; Choi et al., 2016). Our model-based-methodologies (Li et al., 2013b, 2015; Choi et al., 2016; Choi and Francis, 2018) work when utilizing a simulation of the periphery and neural substrate as well. This model based approach, which allows us to directly compared neurophysiological characteristics of VPL stimulation vs. natural touch, or simulated cortical responses to touch (Song et al., 2013; Choi, J. et al., 2015), complements neurosurgical efforts (Hanajima et al., 2004; Ohara et al., 2004; Patel et al., 2006) that have been conducted utilizing electrical stimulation of the Vc thalamus and peripheral nerves directly. Significant work on Vc-MiSt has been conducted during surgical implantation of deep brain stimulators into the VIM thalamus for the treatment of tremor. Stimulation of the Vc in humans produces various paresthesias, especially tactile sensations in the core Vc area. Many MiSt-evoked sensations were reported as tingling, which could be related to the use of high frequency stimulation (about 150 Hz). These stimulus trains were found essential for evoking somatosensory perceptions in many, but not all cases. It has been shown that just a few pulses of stimulation can induce perceptions in humans (Patel et al., 2006).

The possibility of future optogenetic implementations of the basic ideas put forth in this paper have already been shown in the retina (Nirenberg and Pandarinath, 2012). Others have described some of the difficulties with current light sensitive ion channels injected into the somatosensory thalamus (Cruikshank SJ et al., 2010) as well as targeting these into the appropriate areas along with possible solutions to such problems (Yizhar O et al., 2011). A very attractive aspect of these techniques is the fact that the VPL thalamus is a small deep brain structure that can be inject with channelrhodopsins, have them transported to the thalamocortical terminals (Cruikshank SJ,Urabe H,Nurmikko AV and Connors BW, 2010), and then use an optical array at the larger, somatosensory cortex, at least for areas 1 and 2. However, as much of the S1 cortical region related to fine touch and proprioception is buried in the central sulcus in humans and NHPs, it may still be difficult to access without causing some damage. In addition, obtaining fine spatial and temporal optical stimulation at depth below the cortical surface without causing damage is still a challenge. As high-density arrays of micro- and even nano- electrodes are evolving, it is very likely that electrical stimulation will remain the chosen intervention for clinical applications.

